# Climatic clustering and longitudinal analysis with impacts on food, bioenergy, and pandemics

**DOI:** 10.1101/2021.09.30.462568

**Authors:** John Lagergren, Mikaela Cashman, Verónica G. Melesse Vergara, Paul R. Eller, Joao Gabriel Felipe Machado Gazolla, Hari B. Chhetri, Jared Streich, Sharlee Climer, Peter Thornton, Wayne Joubert, Daniel Jacobson

## Abstract

Predicted growth in world population will put unparalleled stress on the need for sustainable energy and global food production, as well as increase the likelihood of future pandemics. In this work, we identify high-resolution environmental zones in the context of a changing climate and predict longitudinal processes relevant to these challenges. We do this using exhaustive vector comparison methods that measure the climatic similarity between all locations on earth at high geospatial resolution. The results are captured as networks, in which edges between geolocations are defined if their historical climates exceed a similarity threshold. We then apply Markov clustering and our novel *Correlation of Correlations* method to the resulting climatic networks, which provides unprecedented agglomerative and longitudinal views of climatic relationships across the globe. The methods performed here resulted in the fastest (9.37 × 10^18^ operations/sec) and one of the largest (168.7 × 10^21^ operations) scientific computations ever performed, with more than 100 quadrillion edges considered for a single climatic network. Correlation and network analysis methods of this kind are widely applicable across computational and predictive biology domains, including systems biology, ecology, carbon cycles, biogeochemistry, and zoonosis research.

## 1 Introduction

With a projected human population of 9 billion by 2050 [1], there is a rapidly increasing demand for food and energy resources. Furthermore, urbanization, human encroachment, and changes in land-use patterns have caused the vast degradation of natural ecosystems and wildlife habitats. This leads to the emergence of multiple zoonotic diseases (diseases caused by microbes or viruses that spread between animals and humans) and thus to an ever increasing frequency of epidemics and pandemics. While *Sustainable Development Goals (SDG) 2030* has given high priority to food security, sustainable agriculture, sustainable energy, climate change, and sustainable use and management of terrestrial ecosystems [2], the implementation thus far has not been successful. Therefore, understanding the patterns related to global climate change and predicting future climate patterns are exceedingly important for the future of sustainable agriculture, bioenergy, and maintainable biodiversity.

Climate is the main driver of organismal adaptation. With a change in climate, species must (i) adapt to the changing climate, (ii) migrate to favorable climatic regions (i.e., range shift), or (iii) face the risk of extirpation from their local habitat [3]. The effects of such climatic changes can be more pronounced in developing countries, where the traditional agricultural practices do not provide enough yield to feed the expanding population. Further, large amounts of agricultural land are under drought stress, and the susceptibility to plant diseases has increased due to altered temperature and precipitation regimes. Thus, the development of climate-resilient, high-yielding, disease resistant crop varieties that can cope with a wide range of environments is of utmost importance. Additionally, lands that are not suitable for agriculture, including marginal lands, can be used for growing bioenergy feedstocks (e.g., lignocellulosic biofuel, an alternative source of sustainable energy). Perennial grasses and fast-growing tree species that can sustain extreme environmental conditions are suitable for this purpose [4]. However, developing high-yielding cultivars/varieties of food and bioenergy crops that are resilient to climate change and pathogens requires careful assessment of global climate patterns.

The shortage of food and energy resources has also compelled human populations (especially in developing countries) to rely on forest resources for food, fodder, and firewood, causing ecosystem imbalance, habitat fragmentation, and loss of biodiversity. High species turnover due to these factors and the changing climate has disrupted the food chain and ecological niches, such environmental stress and habitat fragmentation provides new avenues for viral shedding and zoonotic spillover to human hosts [5]. The major challenge here is to address the need for food and bioenergy while maintaining barriers to zoonotic transmission (i.e., to prevent subsequent pandemics) in the face of climate change.

To address these global crises, understanding the relationships that define climate zones on earth is crucial for future research and planning. Traditionally, climatic analysis has been carried out using the Köppen–Geiger climate classification system [6]. However, it has been suggested that the classical Köppen–Geiger method, with its reliance on heuristic decision rules, should be replaced by a data-driven approach [7]. The primary alternative considered thus far has been the k-means algorithm for clustering environmentally similar geolocations into distinct groups [8]. However, obtaining exact solutions of the k-means problem is NP-hard. Furthermore, the basic k-means algorithm can provably converge to spurious local minima for as few as three clusters [9], and the number of iterations required for convergence is exponential in the number of vectors [10]. Fundamentally, in high dimensions, methods such as k-means and locality sensitive hashing (i.e., methods that are not performing an exhaustive search), are incapable of finding all sets of similar vectors in a data set reliably and efficiently. Furthermore, dimensionality reduction techniques such as Principle Component Analysis (PCA) can lose fine-grained information.

### 1.1 Climate analysis

Our goal here is to enable fine-grained climatic analysis with highly accurate methods that are only achievable by exhaustive similarity search. In this regard, we integrate multiple layers of environmental information to identify climatic zones that share similar environmental relationships around the world and detect how those relationships are changing over time. We take an unbiased, data-driven approach that integrates historical time series observations from 1958 to 2017 of 14 different climate variables extracted from the open-source TerraClimate database [11]. These data are carefully pre-processed to ensure quality control, including cross-variable correlation analysis and distribution normalization (see Section 4.2). We conduct correlation studies using 2-way (all pairs) and 3-way (all triplets) similarity comparisons of equally spaced points of land on earth at high geospatial resolution. Notably, we consider 3-way vector similarities to explore the higher-order interactions of vector triplets compared to 2-way vector pairs, which are traditionally used in correlation studies. We perform (i) a static comparison using an agglomerative 60-year historical view (i.e., 1958-2017) of the environment at all points of land on earth as well as (ii) a longitudinal analysis by considering a series of 10-year time-windows in 1-year step sizes (i.e., 1958-1967, 1959-1968, …, 2008-2017), which results in 51 total time-windows. Specifically, for the 60-year agglomerative view, we use a geospatial grid resolution of 500,710 land coordinates (~19 km^2^), and for the 10-year longitudinal view, we consider 152,100 land coordinates (~35 km^2^).

Due to the massive number of vector comparisons that must be performed (e.g., more than 1.2 × 10^17^ 3-way comparisons for the agglomerative case), we leverage the capabilities of two of the world’s top computing systems. Specifically, we use the Oak Ridge Leadership Computing Facility (OLCF) system, Summit, which is currently the fastest computing system in the United States, and the Jülich Supercomputing Centre (JSC) system, JUWELS Booster, which is currently the fastest computing system in Europe. See S1_Appendix for a detailed description of each. To further increase computational efficiency, we utilize the Combinatorial Metrics (CoMet) library, a data analytics application previously used in comparative genomics studies [12, 13]. The efficiency of CoMet is partly afforded by use of ultralow precision mathematics, meaning that data compression is critical to optimize CoMet performance. Thus, we encode the continuous-valued climate variables that comprise each geolocation into optimal binary representations that (i) retain data context and (ii) maximize performance (see Section 4.2). Finally, to measure similarity between binarized geolocation vectors, we utilize the DUO metric, which is similar to the Sörensen-Dice Index (see Section 4.1). This proportional similarity metric, of which the Sörensen-Dice Index is a special case, has desirable mathematical properties not shared by most other vector similarity measures, hence its attractiveness for this use case [14].

The results of the vector similarity comparisons are captured as networks, in which edges are defined between geolocations if their DUO similarity exceeds a threshold. To identify high-resolution climate zones, we utilize Markov Clustering (MCL), an unsupervised graph clustering algorithm, to define groups of geolocations that share similar environmental relationships. To track how these relationships change over time, we also develop and apply the *Correlation of Correlations* (Cor-Cor) algorithm, a novel methodology that measures similarity between climatic networks corresponding to distinct 10-year time-windows. These methods are used to provide an unprecedented global view of climatic relationships and how they are changing over time (see Section 4.4). An overview of the complete data workflow is shown in Figure 1 and described further in Section 4.

**Figure 1.**
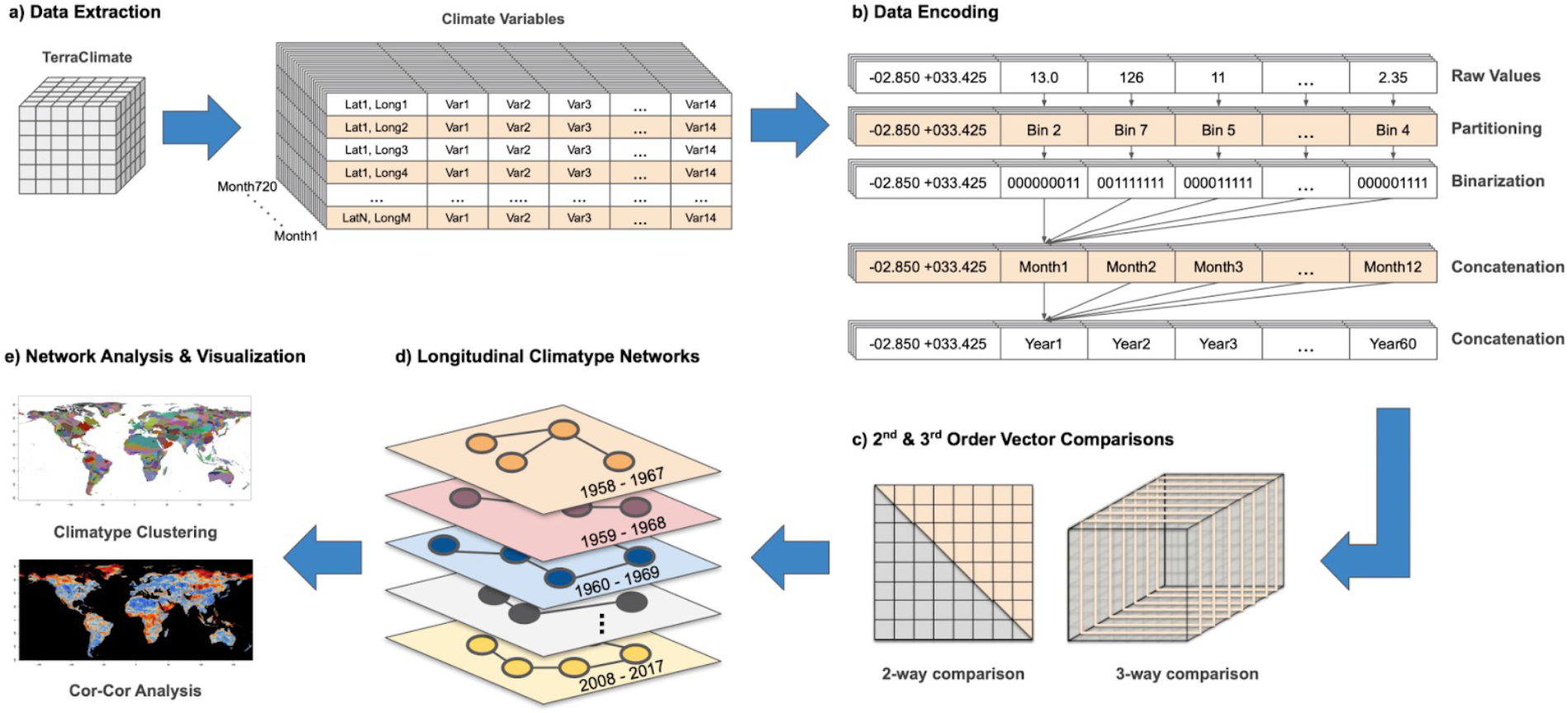
Climate analysis workflow. **a)** Data extraction from the TerraClimate database into geolocation (longitude, latitude) vectors composed of 14 climate layers spanning 720 months of observations. **b)** Vector encoding starts by partitioning raw continuous climate values into discrete bins, followed by binarization using a modified one-hot encoding scheme. The binary representations are concatenated over months and years to produce agglomerative time-windows of historical environmental data. **c)** Exhaustive vector comparisons are performed using highly efficient 2-way and 3-way binary vector comparisons. **d)** Results are translated into network representations of each time-window. **e)** Network analysis and visualization methods are performed, including MCL and the novel Cor-Cor algorithm.

## 2 Results

### 2.1 Scientific results

The scientific results of this work are demonstrated with two viewpoints: (i) climatic clustering and (ii) tracking longitudinal environmental change. Each objective is addressed by formulating a network analysis problem. Climatic networks are constructed by assigning edges between geolocations if their DUO similarity exceeds a threshold. The thresholds are chosen to maintain as many nodes in the network as possible while promoting network sparsity. In particular, the 2-way DUO comparison of the 500,710-coordinate agglomerative data set yields a climatic network with 489,123 nodes and 24,013,802,230 (~24 billion) edges. Similarly, the 3-way DUO comparison applied to the same data set produces a network with 482,987 nodes and 1,620,642,946 (~1.6 billion) edges.

For climatic clustering, in order to cluster the globe according to environmental similarity, we apply the MCL algorithm to the agglomerative climatic networks that span the 60-year range of climate data. The MCL algorithm applied to the 2-way network identifies 830 clusters, while the algorithm with identical tuning parameters applied to the 3-way network reveals 5,238 clusters. A consequence of the different edge counts between 2-way and 3-way climatic networks is seen in the granularity of the resulting clusters, and is visualized in Figure 2. We additionally repeated this analysis for different parameterizations of the MCL algorithm. For climatic clusters between 2-way and 3-way networks across all parameter sets, see https://compsysbio.ornl.gov/.

**Figure 2.**
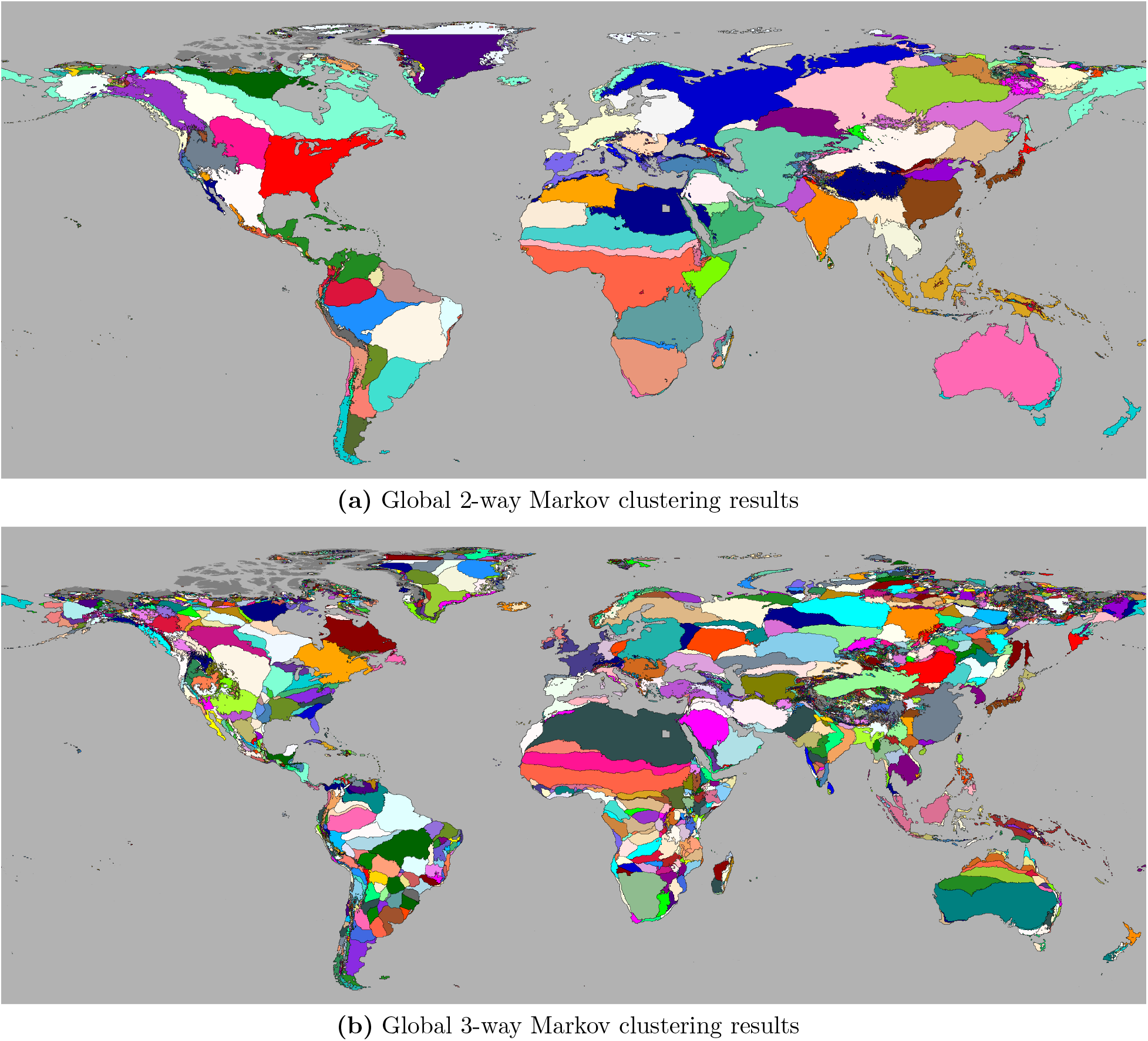
Markov clustering results. Results of applying the MCL algorithm, which indicates high-resolution climatic zones related by environmental similarity. **(a)** 830 climatic clusters arising from 2-way comparisons. **(b)** 5,238 climatic clusters arising from 3-way comparisons.

To track longitudinal environmental change, we apply the Cor-Cor method to the series of 51 10-year time-windows ranging from 1958-1967 to 2008-2017. In particular, we use Cor-Cor to measure the maximum cumulative change over all 10-year time-windows with respect to the first time-window (i.e., 1958-1967), thereby tracking total environmental change over the 60-year period for which there are environmental observations. This analysis was applied to the series of 2-way networks and 3-way networks, and is visualized in Figure 3. We have also produced animations from the Cor-Cor analysis ranging from 1958 to 2017 which highlight global periodicities in the changes detected by the Cor-Cor algorithm over the time series considered here, see https://compsysbio.ornl.gov/.

**Figure 3.**
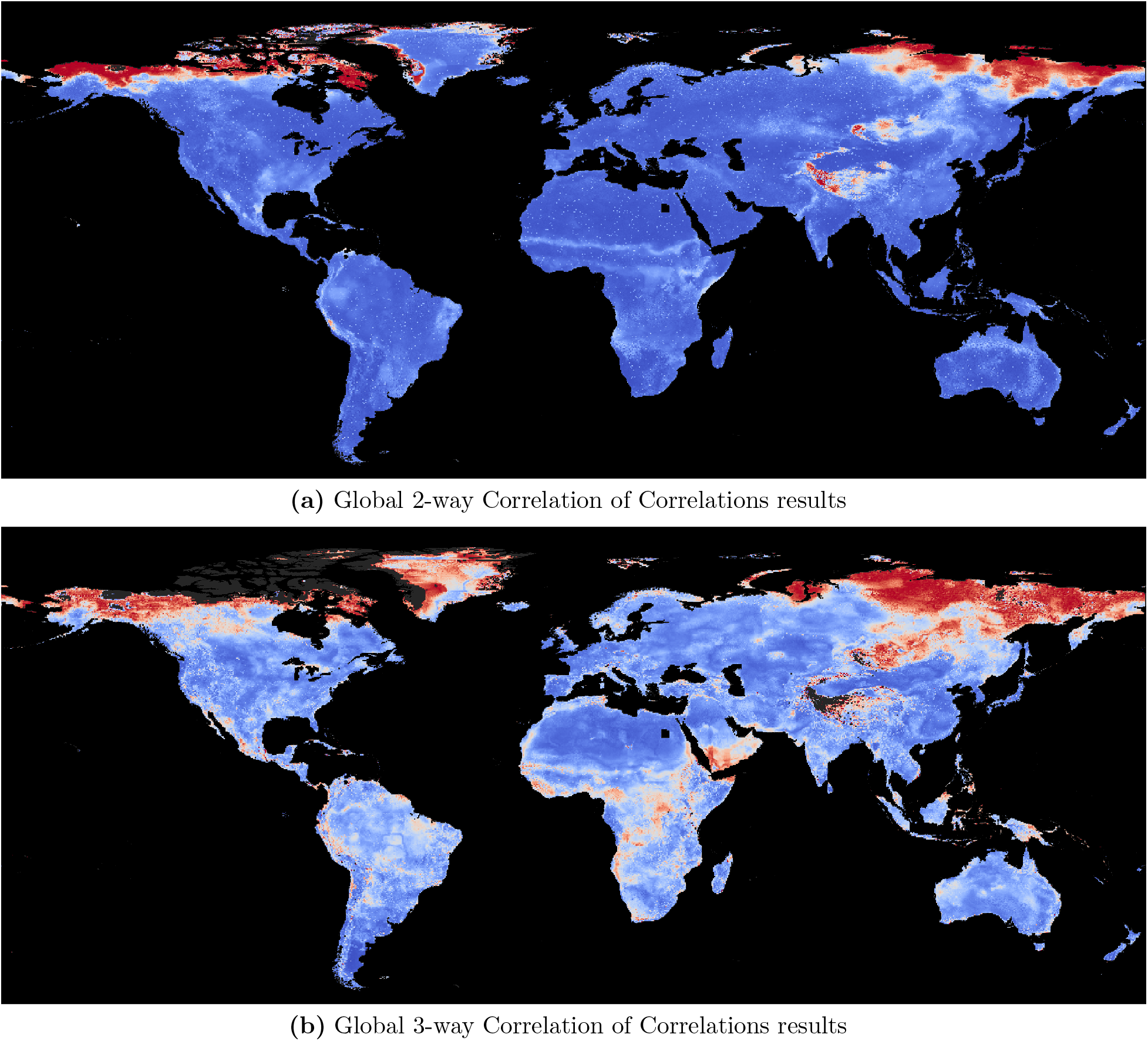
Correlation of Correlations results. Cor-Cor results between the 1958-1967 and 2008-2017 time-windows, indicating areas around the globe experiencing rapid environmental changes. **(a)** shows the Cor-Cor results from 2-way networks and **(b)** shows the Cor-Cor results from 3-way networks. Blue color indicates little-to-no changes in environmental relationships while red indicates large changes.

With the Cor-Cor method using global time series climate data, we are able to highlight regions of earth where the environmental relationships are rapidly changing. These regions may have severe implications for climate change and future zoonotic spillover events. In particular, the Cor-Cor results highlight that the Northern Hemisphere’s high-latitudes are hot-spots for changing environmental relationships, which has consequences related to climate change (see Section 3 for more details). Further, the analysis also reveals potential hot-spots for bat-borne diseases, in particular, Ebola in Africa [15, 16], Hendra virus in Australia [17, 18], and Nipah virus in South and Southeast Asia (especially Bangladesh) [19–21]. The dramatic environmental change in northern latitudes and potential fragmentation of the bat habitats are shown in Figure 4.

**Figure 4.**
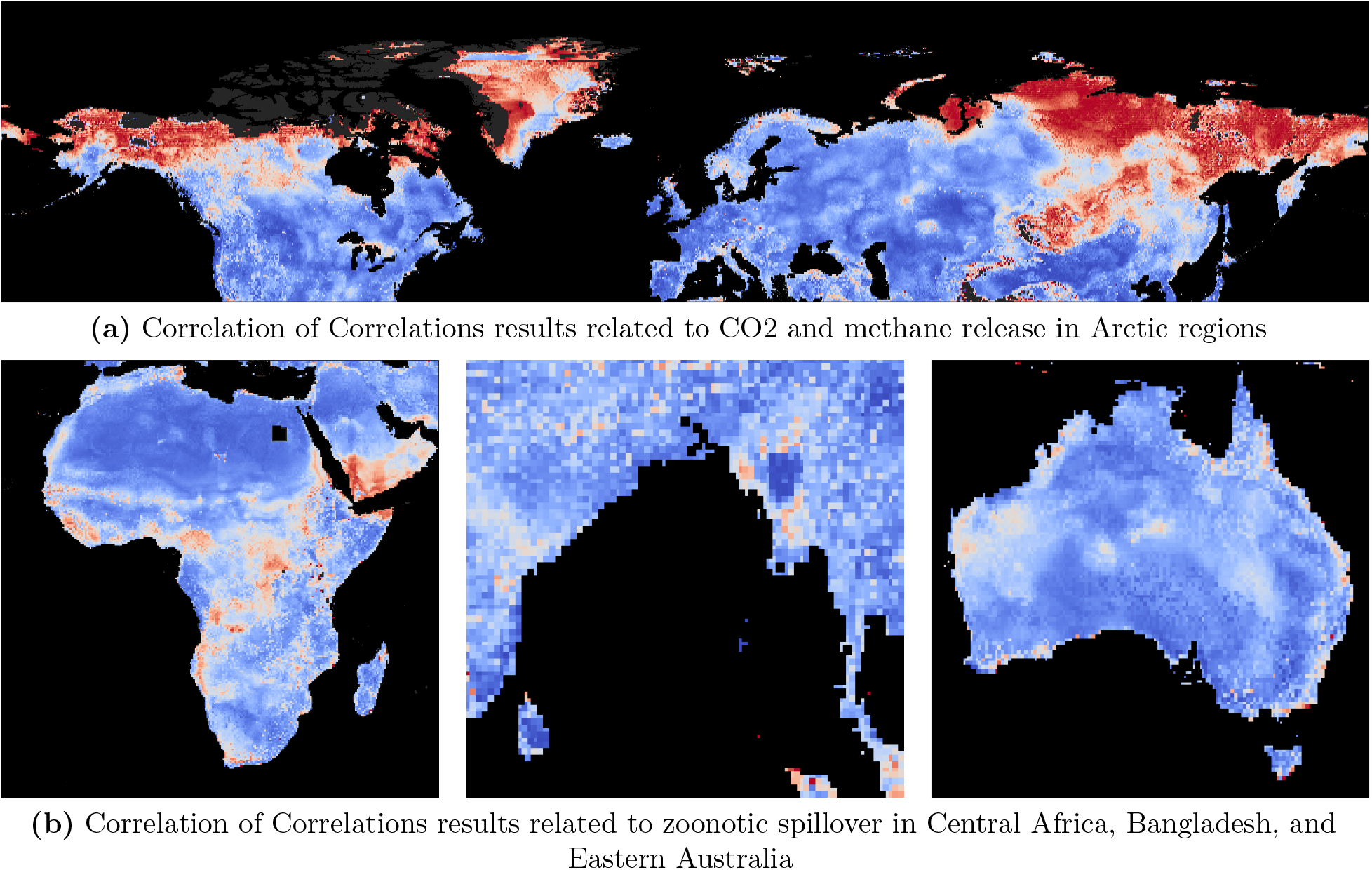
Correlation of Correlations implications. Longitudinal Cor-Cor analysis on 3-way climatic networks between the 1958-1967 and 2008-2017 time-windows reveals rapid environmental changes in **(a)** Arctic regions and **(b)** Central Africa, Bangladesh, and Eastern Australia. Blue color indicates little-to-no changes in environmental relationships while red indicates large changes.

### 2.2 Computational results

Owing to the superior efficiency of computing 3-way comparisons over 2-way, performance data is presented here for 3-way runs using JUWELS Booster. Large DUO comparison runs are split into multiple stages, with each stage writing its own output files that are subsequently combined during post-processing. The 3-way vector comparison was performed on 833 compute nodes of JUWELS Booster, requiring approximately 6.1 hours total wallclock time. The 3-way DUO method was performed in its entirety on the 500,710-coordinate agglomerative dataset with each geolocation vector containing 504,000 binary elements (see Section 4.2 for more information). The computation completed 13 out of 20 stages, after which a node hardware failure occurred. The computation was then restarted to complete the final seven stages. We report on the second part of the computation, for which statistics that were printed at the end of the run were captured.

The operation rate for the core computation was 7.82 × 10^18^ operations per second (ops/sec), and the rate for the entire code execution time was 7.47 × 10^18^ ops/sec. Furthermore, the average rate for the general matrix-matrix multiplication (GEMM) operations alone was 3,866 TeraOps/sec per GPU, 95.4% of the achievable peak 4,050 TeraOps/sec per GPU, and representing GEMM-only performance of 12.88×10^18^ ops/sec across all GPUs used. The entire calculation for the two runs executed a total of 168.7 × 10^21^ operations and ran in 6.1 hours. The equates to an output rate of 585.9 × 10^15^ vector element comparisons per second, 7.21X faster than the figure reported in a similar application [22]. The proportion of time spent for each component of the calculation is shown in Table 1. Each stage is composed of long periods of mostly GPU computation with utilization near 100%, with some overhead for CPU and communication operations. After each of the seven stages, the GPUs are temporarily inactive while output is written. See S1_Fig for GPU utilization over time.

**Table 1.** Computational performance. Timings for execution components during the CoMet vector comparison.

| Component | Time (sec.) | Percent |
| --- | --- | --- |
| Core Computation | 7465.02 | 95.43 |
| Vectors Init | 1.11 | 0.01 |
| Metrics Init | 0.03 | 0.00 |
| Input | 34.49 | 0.44 |
| Output | 321.29 | 4.11 |
| Other | 0.24 | 0.00 |
| TOTAL | 7822.17 | 100.00 |

The algorithms used here exhaustively compute all 2-way or 3-way comparisons, but only write a small fraction of significant results to output files based on a given DUO similarity threshold (i.e., 0.70 for 2-way and 0.55 for 3-way DUO). The corresponding number and size of input and output files for the four cases considered here are shown in Table 2.

**Table 2.**
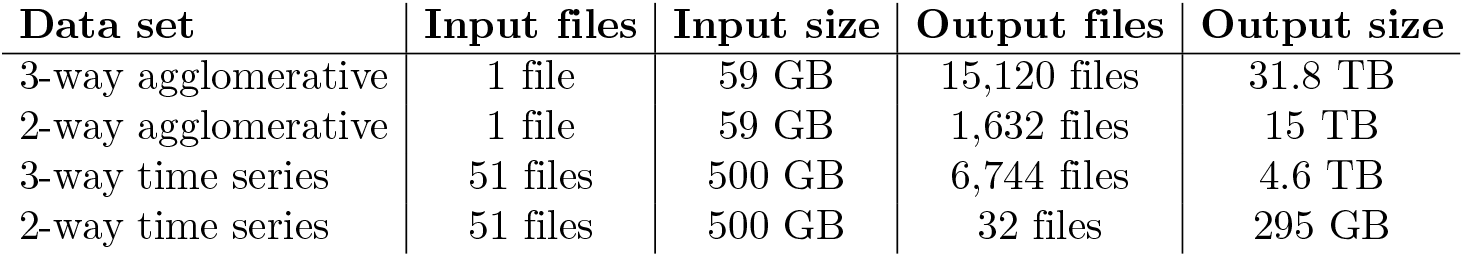
Data set characteristics. Number and size of input and output files for 2-way and 3-way vector comparisons of the agglomerative 60-year climate data set and the series of 51 10-year time windows.

| Data set | Input files | Input size | Output files | Output size |
| --- | --- | --- | --- | --- |
| 3-way agglomerative | 1 file | 59 GB | 15,120 files | 31.8 TB |
| 2-way agglomerative | 1 file | 59 GB | 1,632 files | 15 TB |
| 3-way time series | 51 files | 500 GB | 6,744 files | 4.6 TB |
| 2-way time series | 51 files | 500 GB | 32 files | 295 GB |

### 2.3 CoMet scaling

To measure scaling on JUWELS Booster, a weak scaling study was conducted using artificial data derived from a realistic climate data set containing 500,710 vectors of length 1,008,000 elements each, corresponding to full coverage of the earth’s land mass (excluding Antarctica) at ~ 19 km^2^ resolution. The 3-way DUO comparisons were computed at up to 825 compute nodes of JUWELS Booster. For benchmarking purposes, we only consider one of five compute stages, with each run taking approximately two hours of wallclock time. At the largest node count, the CoMet core computation achieved 9.37 × 10^18^ operations per second (ops/sec), and the entire application, including I/O, ran at 9.21 × 10^18^ ops/sec. The core computation GEMM rate is 3.97X the rate of 2.36 × 10^18^, which was previously reported [22]. The average GEMM-only performance was 3,985 ops/sec, very near to the 4,050 achievable peak for the A100 GPU (i.e., the GPU used on the JUWELS Booster computing system). Individual GEMMs have the form *C* = *A^T^ B* where *A* and *B* have dimensions roughly 1, 008,000 × 16, 704, each performed in approximately 0.14 seconds on an A100 GPU. See S2_Fig for a visualization of the weak scaling performance on JUWELS Booster.

To measure performance on Summit, we solved the 2-way DUO correlation problem using an additional climate data set with 8,834,910 vectors of length 504,000, corresponding to a resolution of ~4 km^2^. Here we achieved 2.09 × 10^18^ GEMM operations for the core algorithm calculation on 4,624 nodes of Summit, equaling 66.2% of achievable mixed precision peak on Summit. The mixed precision FP16/FP32 GEMMs alone ran at 104.65 TeraOps, or 92.6% of the achievable peak. The entire core calculation required 75 seconds to execute on Summit.

Note that the data sets used for performance studies are not used for downstream network analysis since they differ in number and length of geolocation vectors. In particular, we utilize the 152,100-coordinate and 500,710-coordinate data sets to compare differences between the 2-way and 3-way climatic networks using identical data sets and algorithm parameters.

## 3 Discussion

The use of 3-way vector comparisons to generate climatic networks has produced significant scientific insights into the time-dependent environmental processes and shifts around the globe. These improvements are directly related to the features of the resulting climatic network. An important observation is the trade-off between computational burdens. Clustering the 2-way network, with more than 24 billion edges, is significantly more computationally expensive compared to the 3-way network with just over 1.6 billion edges. However, computing a 3-way vector comparison is orders of magnitude more expensive compared to computing 2-way vector similarities. Yet, the increased computational burden of the 3-way vector comparison is offloaded to the extremely efficient CoMet library, which enables broader scientific use and analysis of the emergent graphs. The differences between 2-way and 3-way comparisons are also highlighted in the MCL and Cor-Cor results, in which the 3-way climatic networks reveal rapidly changing regions with significantly higher sensitivity and granularity compared to the 2-way graphs, as shown in Figure 2 and Figure 3. These results suggest that the 3-way DUO metric captures more higher-order interactions among geolocations, thereby yielding networks that are richer in information content (i.e., have a higher signal/noise ratio), and, thus, are better data-driven models of the global environment.

The MCL results shown in Figure 2 highlight different regions around the globe that share similar climates. Importantly, the clusters naturally form into geospatially auto-correlated groups, despite the observation that the DUO vector comparisons and MCL algorithm are not given coordinate-level information (e.g., latitude, longitude, elevation, etc.). This phenomenon emerges from the inherent auto-correlation structure of the climate data, in which adjacent points of land share similar environmental features. Thus, the global clusters exhibit expected grouping in both the 2-way and 3-way cases. In particular, cluster granularity increases as a function of elevation and climatic extremes. For example, in mountainous regions, small changes in latitude or longitude can have dramatically different climates, which results in a larger number of small clusters (e.g., the Himalayan mountain range in Central Asia). In contrast, other areas with more consistent climates are captured as larger clusters that can span continents (e.g., the Sahara Desert in Northern Africa).

The Cor-Cor results highlight that the Northern Hemisphere’s high-latitudes are hot-spots for environmental change, as seen in the top of Figure 4. It is reported that 25% of the world’s soil carbon is stored as recalcitrant peat [23]. Thus, as the permafrost melts, these frozen stores of recalcitrant peat are being exposed to microbiotic conversion (via methanogenic bacteria) from peat to methane and CO2. Of note is that methane has a 21-fold higher global warming potential than that of CO2 [24]. As such, rapid environmental changes in these areas have the potential to significantly impact global carbon fluxes, and therefore further accelerate climatic changes [25, 26].

The Cor-Cor results also align with the hypothesis that changes in land-use patterns cause fissioning of bat populations, leading to nutritional stress, which in turn facilitates viral shedding and spillover to human and livestock hosts [27], as seen in the bottom of Figure 4. The high-resolution time series data, together with other data sources such as genomic, phenotypic, societal and behavioral data, can have important implications for a better understanding of the zoonotic reservoir host and pathogen biology, and also for detecting the factors driving the cross-species transmission.

Broadly speaking, the analyses of these networks allows the scientific examination of the relationships between micro-, mezo-, and macro-climates, and the processes that they drive, including carbon sequestration/release as greenhouse gases, glacier/ice retreat, sea-level rise, sustainable agriculture, and changes in taxonomic composition and ecosystem stability.

### 3.1 Computational innovations

This work takes advantage of the continuing trend of GPU accelerators to deploy more hardware features to increase processing rates on increasingly large workloads. The Nvidia Ampere GPU computes matrix products on 64 individual 1-bit inputs at a rate 4X faster than one 64-bit double precision input using tensor cores, and 8X faster than using the standard double precision compute units. Our previous methods employed FP16/FP32 arithmetic, which runs 16X slower than 1-bit GEMMs on the Ampere architecture; without the new 1-bit GEMM capability, the runs completed in this work would require the same number of GEMM operations, however would run roughly 16X slower. The use of lower precision capabilities enables substantially higher throughput and accuracy from the use of larger models as well as greater power efficiency and cost reduction for these computations.

The 6-hour DUO calculation documented here executed a total of 168.7 low precision ZettaOps, or 1,952 low precision PetaOp/s-days. This is in the same magnitude range as training the massive GPT-3/large language model, which required 314 mixed precision ZettaOps [28] at estimated training cost of $4.6 million [29], and AlphaGo Zero, which required 1,860 Petaflop/s-days [30]. These are some of the largest calculations ever completed to date. It is suggested that within a few years, a billion dollars could be spent on training a single deep learning model [31]. It is clear that heterogeneous hardware features must be exploited in every way possible to support these large-scale projects.

Processor heterogeneity is driven in part by the enormous requirements of deep learning workloads as well as the post-Moore’s Law era of computing. Reduced precision arithmetic is now supported on processors from all major HPC processor vendors, including Nvidia, AMD, and Intel; these should be directly usable by the methods described in this work. Upcoming HPC systems such as NERSC’s Perlmutter and CINECA’s Leonardo will also use Ampere GPUs and support the 1-bit GEMM methods described here.

Mixed precision computing has gained significant interest in HPC in recent years. One lesson learned from this work is the need to optimize all parts of the application workflow to keep up with the performance boost from reduced precision, from both compute and data transfer speed improvements. Accelerating GEMM operations by 16X suddenly exposes other parts of application runtime that must be optimized. Additionally, careful attention must be paid to accuracy and algorithmic stability issues. The accuracy of the algorithms treated here is not impacted by lower precision methods, however some algorithm classes require careful numerical analysis considerations to ensure reliability.

The optimization of data motion and management of data within the workflow are critical. Had the output data not been filtered, the 6-hour simulation described here would have written 2.68 exabytes of output data; the actual figure after filtering by threshold was 4.85 TB. Important data-related themes we encountered in this work include: filtering the data as early as possible after creation; considering how to perform more *in situ* streaming analytics before filtering; bringing the compute closer to the data, for example, by moving more of the computation to the GPU; optimizing algorithm data motion to keep up with radically increased processor speed; optimizing processing and transfer costs for the most expensive compute resource and offloading pre- and post-processing to other resources; and organizing data/compute workflow processes and tools to manage the data pipeline.

Data analytics methods and machine learning techniques are playing an increasing role in HPC workloads. The word “data” does not occur in the text of the 2004 High-End Computing Revitalization Act [32], however, is clearly visible in the 2015 executive order creating the National Strategic Computing Initiative [33]. Scientific discovery at large scale using data-driven methods is becoming more common, as evidenced by the recent AlphaFold2 results and SARS-CoV-2 spike dynamics findings [34]. Interest is also growing in coupling large computing resources to devices at experimental facilities.

The methods developed here are applicable to many other problem domains for which it is required to identify networks of similarity relationships between elements in large quantities of data, representable by vector similarity measures. These problem domains include systems biology as well as ecology, materials science, carbon cycles, biogeochemistry, additive manufacturing, and zoonosis research, to name a few.

### 3.2 Conclusions

Traditional climatic analyses have relied on the use of heuristics and climate models to draw scientific conclusions about global climate patterns. A major contribution of this work is our unbiased, model-free, data-driven approach, which minimizes model discrepancy and helps ensure the objectivity of the results herein. Further, each climate variable is carefully pre-processed to reduce cross-variable correlations, account for extreme outliers, and ensure an even variable distribution for the DUO similarity metric. Taking these pre-processed variables together with exhaustive 2-way and 3-way all-against-all vector comparisons produces extreme-scale climatic networks that are comprised of deeply complex environmental relationships. We exploit these relationships using (i) MCL to divide the globe into environmentally similar high-resolution zones, and (ii) our novel Cor-Cor method, which tracks environmental changes over time across each location on the planet. The conclusions drawn from this work affect a number of critical scientific use cases ranging from sustainable bioenergy and food production to detecting the areas of the world most susceptible to zoonotic spillover events and subsequent deadly pandemics.

In contrast to typical dense vector similarity calculations, which compare all vector pairs in a given data set, we consider a 3-way vector similarity calculation between all vector triplets. This comparison reveals higher-order interactions that are otherwise missed using pairwise comparisons. Further, the thresholded climatic networks that emerge from the 3-way DUO comparison are more sparse (i.e., include fewer edges) compared to the 2-way comparison, which reveals more granular and scientifically relevant features related to climatic regions.

A primary aspect of this work is the analysis of time-dependent trends across the earth using a sliding 10-year window. The addition of a longitudinal dimension by which we can reason about temporal processes is a crucial step in the analysis of historical environmental patterns as well as making projections about future climate trends. Enabled by such longitudinal climatic networks, this work introduces Cor-Cor, a novel method for measuring environmental change at every location across the globe by directly comparing climatic networks throughout history. This allows scientists to track shifts in complex environmental relationships at global scale to identify which areas on earth are experiencing the most rapid changes. These analyses have important implications on sustainable agriculture (food and bioenergy) and zoonotic spillover events that can lead to pandemics, and thus multiple impacts on human health and well-being.

Additionally, the computational innovations developed in this work are aligned with the confluence of two trends in scientific computing. On one hand, observational, experimental, and simulation data are increasingly fueling scientific discovery, with data volumes growing at a rate that outstrips available compute capacity, thereby increasing the need to bring the compute closer to the data. On the other hand, the “end of Moore’s law” theme is leading to new approaches, custom processors, and heterogeneous on-die compute units to continue the growth in computing capacity. These changes necessitate the relentless optimization of codes in order to keep pace with the shifting balances of performance rates for the different compute components. To meet these challenges, the CoMet application was developed as a data analytics code for unsupervised clustering based on vector similarity methods for large volumes of data [12, 13, 22]. Alternative vector similarity methods based on locality-sensitive hashing [35] are unable to find all important similarities between vectors in an efficient way. Such methods suffer from curse of dimensionality issues, and thus costs grow exponentially with required accuracy. Thus, CoMet enables accurate vector similarity clustering through complete, exhaustive search.

The work presented here makes use of publicly available data and software. In particular, we utilized the open source TerraClimate data repository [11] for historical climate data, the CoMet software library (see https://github.com/wdj/comet) to conduct 2-way and 3-way vector comparisons, and HipMCL for large-scale unsupervised climatic clustering [36].

## 4 Methods

### 4.1 DUO similarity metric

The large number of computations required by the exhaustive similarity search necessitates the use of high-performance computing (HPC) systems. Thus, in this work, we leverage the capabilities of two of the world’s top computing systems to scale 2-way and 3-way global climate vector comparisons: Summit and JUWELS Booster. Additionally, we use the CoMet library, which takes advantage of ultralow precision mathematics (e.g., 1-bit GEMMs, which are roughly 16X faster than FP16/FP32), and requires that the geolocation vectors are converted to binary. To measure similarity between binary geolocation vectors, we consider the DUO metric, now implemented in CoMet. The DUO metric resembles the Sörensen-Dice Index of vector similarity (a statistic for comparing discrete distributions), however, it has been adapted to compute multiple correlation statistics between vectors. For computational efficiency, DUO calculates the similarity between two binary vectors by binning matrix values into low or high (0 or 1) categories [37–40]. DUO comparisons produce four possible combinations for a 2-way comparison: (1, 1), (1, 0), (0, 1), and (0, 0). The DUO metric between binary vectors *i* and j is given by the equation:

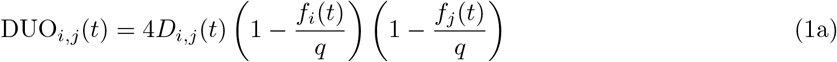

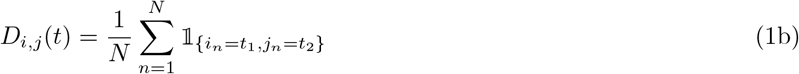

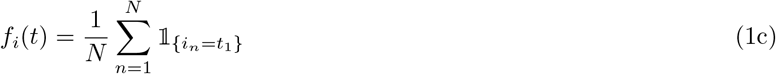

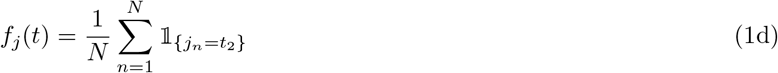

where *t* = {(*t*_1_, *t*_2_) | *t*_1_, *t*_2_ ∈ {0,1}} represents one of the four possible combinations, *q* ∈ [1, ∞) is a scaling factor, 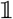 represents the indicator function (e.g., 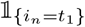 is equal to 1 when the *n*^th^ element of vector *i* is equal to *t*_1_ and 0 otherwise), and *N* is the total length of the geolocation vectors. The term *D_i,j_* (*t*) represents the proportion of elements exhibiting relationship *t*, while *f_i_*(*t*) and (*t*) model the probability of observing the corresponding binary element (*t*_1_ or *t*_2_) in vector *i* and *j*, respectively. In other words, *D_i,j_*(*t*) is equal to the fraction of vector lengths of when *t*_1_ and *t*_2_ co-occur, and *f_i_*(*t*) and *f_j_* (*t*) are the fraction of vector elements in *i* and *j* that are equal to *t*_1_ and *t*_2_, respectively. This metric calculates the correlation (and effectively anti-correlation) values between all compared vectors. For example, a region of high values in vector *i* and high values in the same region in vector *j* could correlate. Conversely, a region of high values in vector *i* and low values in the same region in vector *j* could also correlate. Additionally, DUO accounts for frequency effects by scaling the resulting values according to the fraction of high/low values in vectors being compared. Following [37], we set the scaling factor, *q*, equal to 1.5.

The DUO metric is also extended to 3-way vector comparisons, which result in eight possible comparison combinations (i.e., (1, 1, 1), (1, 1, 0), (1, 0, 1), …, (0, 0, 0)). The 3-way DUO metric for binary vectors *i*, *j*, and *k* is given by the equation:

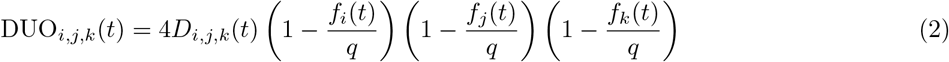

with *t* = {(*t*_1_, *t*_2_, *t*_3_) | *t*_1_, *t*_2_, *t*_3_ ∈ {0,1}} and the terms *D_i,j,k_*(*t*), *f_i_*(*t*), *f_j_*(*t*), *f_k_*(*t*), and *q* following similarly from Eqs (1a–1d). Notably, vector pairs/triplets with DUO scores approaching 1 indicate environmental similarity, and those with DUO scores near 0 are considered dissimilar.

### 4.2 Data generation and pre-processing

The climate data we consider in this work are extracted from the TerraClimate database, which contains 14 layers of continuous-valued climate data: minimum temperature, maximum temperature, vapor pressure, precipitation accumulation, downward surface shortwave radiation, wind-speed, reference evapotranspiration (ASCE Penman-Montieth), runoff, actual evapotranspiration, climate water deficit, soil moisture, snow water equivalent, and the Palmer drought severity index [11]. Each climate layer spans 62 years, corresponding to 744 monthly observations, and ranges from January, 1958 to December, 2019 between −90 and +90 latitude and −180 and +180 longitude.

In this work, climate data is extracted at two geospatial grid resolutions: 152,100 land coordinates (~35 km^2^), and 500,710 land coordinates (~19 km^2^). See S3_Fig for an example coordinate grid over the contiguous United States. Geolocations corresponding to bodies of water and Antarctica are not considered in this work (i.e., we exclude geolocations between −90 and −60 latitude). For each grid coordinate, we extract the corresponding climate data using the `xarray` software package in Python, which utilizes a geometric transformation to convert between world and pixel coordinates, and utilizes a nearest neighbor search to locate corresponding values of a given climate layer [41]. Further, we take additional steps to ensure that grid coordinates are never sampled twice and we discard any coordinates with missing observations. Using this strategy, for each geolocation in the 500,710-coordinate set, we construct agglomerative data vectors that span the full time range of observations. For the 152,100-coordinate data set, we construct a series of data vectors composed of 10-year time-windows in 1-year increments. The agglomerative data set is used to provide a full view of correlated geographic locations while the time-windows are used for a longitudinal study that reveals how climatic relationships are changing over time.

Multiple data pre-processing steps are taken to ensure quality control, including cross-variable correlation analysis, distribution-agnostic variable partitioning, and conversion to binary. To test for correlations between climate layers, we performed a pairwise correlation analysis using Pearson’s correlation coefficient across all 62 years of data (the total number of years of data currently available in the TerraClimate database) over the 500,710-coordinate grid. This analysis revealed strong correlations between the minimum and maximum temperature (e.g., median Pearson’s correlation of 0.97 across all 62 years). Thus, to avoid duplicated information arising from strong correlations, we derived a new climate layer: temperature range (i.e., the difference between maximum and minimum temperature) to replace the maximum temperature layer. In this way, we retain the same level of information (i.e., maximum temperature can be derived from temperature range and minimum temperature), but the median correlation was reduced to 0.373. Further, the correlation analysis revealed that the final two years of data (i.e., 2018 and 2019) contained layers that were highly skewed compared to the rest of the climate variables. See S4_Fig for a snapshot of the pairwise correlations. Therefore, we removed these two years of data from all climate layers and analyses to avoid biases that may have been introduced. Thus the total time range is reduced to 60 years, or 720 monthly observations, meaning that for the longitudinal study, we consider 51 time-windows (i.e., 1958-1967, 1959-1968, …, 2008-2017).

In order to carry out an efficient large-scale 1-bit GEMM vector comparison, a precise variable encoding scheme (i.e., conversion from continuous-valued to binary-valued data) is essential. Encoding begins with the partitioning of the raw continuous climate variables into discrete bins. Due to the wide range of distributions and scales that span the climate layers, we apply an equal-frequency binning methodology (similar to a uniform quantile transform) to standardize each layer. This process transforms the distribution of each climate layer into a uniform distribution and seamlessly accounts for extreme outliers.

To balance computational efficiency with scientific accuracy, we encode each bin into a binary representation with 50 bits. The binary encoding scheme is a modified one-hot encoding, where, for a binary representation with *n* bits and a given bin, *k*, the binary assignment is 0^*n–k*^ 1^*k*^. For example, for bins *k* = 1, 2,…, 5 using *n* = 5 bits, the binary representations are 00001, 00011, 00111, 01111, and 11111, respectively. This encoding scheme is harmonious with the DUO metric since the high and low values of climate layers between geolocations are aligned. For example, consider adjacent bins 3 and 4 of a climate layer. Since the bins are close to one another, they should correlate more strongly. In our encoding scheme (with *n* = 5 for simplicity), the corresponding binary representations are 00111 and 01111, which correlate since the two representations differ by only one bit. Note that the bit encoding approach used here could be converted into a method that more directly operates on floating-point or integer arithmetic, as described in [14]. However, such methods would require an absolute value or scalar minimum operation, and since these are not implemented in GPU tensor core hardware, such methods would run at significantly slower speeds. Finally, to create agglomerative time-windows for each geolocation, the binary representations of the 14 climate layers are concatenated over months and years. This produces binary vectors of length 504,000 for the 60-year view (i.e., 14 layers × 12 months × 60 years × 50 bits), and similarly 84,000 for the 10-year time-window view.

### 4.3 Vector comparison

Using the binarized climate data for both the agglomerative 60-year data set with 500,710 geolocations and the longitudinal series of 51 10-year time-windows with 152,100 geolocations, we perform both 2-way and 3-way vector comparisons using the DUO vector similarity metric implemented in CoMet. In the 2-way comparison, the similarity between all unique pairs of geolocations are measured using Eq 1a. Similarly, for the 3-way comparison, all unique triplets of geolocations are compared using Eq 2. The total number of 2-way and 3-way comparisons for *N* vectors is *N*^2^ and *N*^3^, respectively. Thus, the 152,100-coordinate produces more than 2.3 × 10^11^ 2-way and 3.5 × 10^15^ 3-way comparisons. Similarly, the 500,710-coordinate data set results in more than 2.5 × 10^11^ 2-way and 1.2 × 10^17^ 3-way comparisons. The extreme scale and computing requirements of such numbers requires multiple computational innovations.

Previous versions of CoMet embodied multiple innovations to adapt the targeted methods to modern leadership-class systems [22], enabling up to a five orders of magnitude improvement over the previous state of the art, including multidimensional hierarchical parallelism, use of GPU accelerators, reformulation of the targeted algorithms to use high performance GEMM operations, asynchronous overlap of compute and transfers, load balancing methods, and use of reduced precision compute units (e.g., tensor cores). However, solving the extreme-scale problems addressed in this work requires further innovations that also align with expected future hardware trends.

Largely motivated by requirements for training deep learning (DL) models, recent GPUs feature special-purpose compute units for matrix product operations (e.g., Nvidia GPU tensor cores and AMD MI-series GPU matrix core engines [42]). Most recently, Nvidia Turing and Ampere architectures offer hardware for computing GEMM matrix-matrix products on ultralow-precision 1-bit inputs accumulated to INT32 outputs. These are already being exploited for some computational problems [43, 44]. The Turing architecture offers a non-conventional XOR-based matrix product that we have adapted to computing the standard GEMM result in CoMet. The Ampere architecture additionally offers a standard 1-bit GEMM that is usable by CoMet directly. The 1-bit GEMMs offer theoretical peak performance 16X faster than the standard half-precision GEMMs, and 256X faster than double-precision matrix product performance [45]. The DUO algorithm takes 1-bit indicator values as inputs, making it directly amenable to this kind of computation. Since these GEMMs are computed, accumulated, and stored as full INT32 values, exact bit-for-bit equivalence with higher precision implementations is maintained.

The extreme speed of 1-bit GEMMs must additionally be matched by increasing the effective transfer speeds in other parts of the computation. To achieve this, the process used to discard thresholded connections between vectors (i.e., pairs/triplets of dissimilar geolocations) is reimplemented on the GPU, and the values not thresholded are losslessly compressed before returning to the CPU by using the Nvidia CUB library. This makes the implementation more throughput-friendly and frees up large amounts of CPU memory, enabling much longer asynchronous pipeline depths and lower overheads. Additionally, other calculations such as matrix formation have been moved to the GPU, resulting in faster speed and less CPU-GPU communication.

The original CoMet communication pattern, identical to the mpiGraph benchmark [46], required every rank to communicate with every other rank at some point in the computation. For the communication-intensive 2-way vector comparisons, this has been changed to a simpler persistent ring (circular shift) pattern, which is more embeddable into common interconnects such as dragonfly topologies with less risk of network congestion. Additionally, GPU-aware MPI has been deployed for use with systems that benefit from it. Further, the original 3-way comparison algorithm described in [22] and [12] was based on computing 3 GEMMs to populate the 8 possible comparison values for each vector triplet. An algorithmic modification has been made requiring only 2 GEMMs, each of these computing 4 of the 8 relevant comparisons for the three geolocation vectors. This method gives results identical to the previous method but increases the throughput rate by up to 50%.

We use the methodology described in [22] to measure computational performance. Timings are collected for parts of the computation with the `gettimeofday` function, using `cudaDeviceSynchronize` and `MPI_Barrier` to synchronize nodes. This is done infrequently to avoid unnecessarily interrupting asynchronous operations. Unless otherwise specified, the core algorithm computation time is measured without I/O. Data input time is typically a small fraction of runtime. Output time is highly dependent on the chosen output threshold factor, however this setting is calibrated to minimize the impact of output time cost while maintaining scientific fidelity. Operation counts are accumulated manually in the code during execution. The standard definition of a GEMM operation is used: for the product *C* = *AB* and matrices *A* and *B* of dimensions *m* × *k* and *k* × *n*, the GEMM operation count is 2*mnk*, for the given precisions of the matrix entries of *A, B*, and *C*. GEMM-only results are reported based on this operation count. For the core metrics computation and vector comparisons, only operations meaningful to the scientific results are counted; for example, calculations on matrix size padding and required computations of unused values of the result matrix are not counted as part of the total.

Finally, in order to complete the vector comparisons on the encoded data sets with CoMet, the input files were transferred from OLCF to JSC. After performing the 3-way global climate vector comparisons on JUWELS Booster, the resulting output data was then transferred back to OLCF for post-processing. Due to the large amount of data that needed to be transferred, we leveraged the multi-stream capabilities of UFTP to transfer the entire CoMet output from JSC with a transfer rate of over 140MB/s using eight streams. For example, UFTP allowed a complete data transfer for the 4.6TB data set in approximately 10 hours. Without a multi-stream capable tool, the same transfer would have taken more than 81 hours using `scp` or `rsync`, which could only achieve a bandwidth of approximately 17MB/s.

### 4.4 Network analysis

In order to construct climatic networks from the set of DUO vector comparisons (where each vector corresponds to a point of land on earth), vector pairs/triplets whose DUO similarity exceeds a given threshold are stored in a graph as nodes (i.e., geolocations) with edges linking the pair/triplet. An edge is created for each vector pair passing the threshold, while three edges are created for each vector triplet. Further, we use the DUO similarity scores as edge weights. By using a threshold to retain edges between geolocations with high DUO similarity scores, a sparse network structure emerges that is then used for downstream scientific analysis.

Since the results from large numbers of 2-way and 3-way combinations can quickly lead to writing exabytes of data, care must be taken in choosing an appropriate threshold that balances network sparsity with output size. Thus, we apply a sub-sampling scheme inspired by sparse grids to quickly test thresholds and resulting output sizes in order to extrapolate to the full vector comparison. See S5_Fig for as example sparse grid sample over the contiguous United States. The sub-sampling scheme is chosen to reduce the total number of geolocations while also preserving the auto-correlation structure of nearby geolocations (i.e., adjacent locations on earth have similar climates). By computing comparisons over the sub-sample of geolocation vectors, we approximate the DUO similarity distributions over all points on earth and define which proportion of DUO correlations are saved, thereby defining the size of the data that will be stored. Through this experimentation, the thresholds are determined to be 0.70 for 2-way DUO and 0.55 for 3-way DUO for the data sets considered here.

To identify high-resolution climate zones, we apply Markov Clustering (MCL), an unsupervised graph clustering algorithm, to the 500,710-coordinate agglomerative 60-year climatic networks that emerge from CoMet using the `HipMCL` [36] high-performance clustering package on the Summit supercomputer at OLCF. MCL is an iterative method that groups nodes into clusters based on the graph topology. Since nodes in a climatic network represent geolocations and edges represent environmental similarity, the clusters derived from MCL highlight areas around the globe that share similar climatic features. We use MCL in this work since, unlike k-means, the number of clusters is not assumed to be known *a priori*. Further, the granulatity of the clusters can be controlled by adjusting parameters of the MCL algorithm (e.g., inflation rate). For the results shown here, we set the inflation rate to 4.0. The results for inflation rates 2.0, 4.0, and 6.0 can be seen at https://compsysbio.ornl.gov/.

Next, we conduct a longitudinal analysis of the 152,100 coordinate 10-year time-window networks by developing and applying the Cor-Cor algorithm, a novel methodology to track environmental changes at every geolocation across the globe. Formally, the Cor-Cor algorithm uses the Sörensen-Dice Index to measure changes in a node’s adjacency vector between two corresponding climatic networks. This comparison is possible because the nodes (i.e., geolocations) between any two climatic networks are the same, however, the neighborhoods of each node (as defined by the edge set) can vary depending on the environmental relationships of each location to all other locations on earth. In this way, areas with Sörensen-Dice scores near 1 indicate little-to-no change in environmental relationships, whereas areas with scores near 0 indicate large changes. Applying this methodology to the series of longitudinal climatic networks reveals which areas around the globe are experiencing the most rapid environmental changes.

## 5 Acknowledgments

The authors would like to thank Bronson Messer and Don Maxwell of the OLCF; Thomas Lippert, Norbert Attig, Damian Alvarez, Wolfgang Frings, Andre Giesler, Andreas Herten, and Benedikt von St. Vieth of the Jülich Supercomputing Center for access and assistance with JUWELS Booster; Brandon Cook, Muaaz Awan, and Paul Lin of NERSC for access and assistance with the Cori-GPU DGX/A100 system; and Jack Wells, Axel Koehler, Jiqun Tu, and Max Katz of Nvidia. This research used resources of the Oak Ridge Leadership Computing Facility allocated by the DOE ALCC program and was supported by the Center for Bioenergy Innovation, the Plant-Microbe Interface SFA, the Feedstock Genomics Program, the Exascale & Petascale Networks for KBase project, the Integrated Pennycress Resilience Project (all supported by the Office of Biological and Environmental Research in the DOE Office of Science). Funding was also provided by the United States Government as well as the National Institute on Aging of the National Institutes of Health under project 1RF1AG053303-01. This research used resources of the National Energy Research Scientific Computing Center (NERSC), a U.S. Department of Energy Office of 638 Science User Facility located at Lawrence Berkeley National Laboratory, operated under 639 Contract No. DE-AC02-05CH11231. The manuscript was authored by UT-Battelle, LLC under Contract No. DE-AC05-00OR22725 with the US Department of Energy. The US Government retains and the publisher, by accepting the article for publication, acknowledges that the US Government retains a nonexclusive, paid-up, irrevocable, worldwide license to publish or reproduce the published form of this manuscript, or allow others to do so, for US Government purposes. The Department of Energy will provide public access to these results of federally sponsored research in accordance with the DOE Public Access Plan (http://energy.gov/downloads/doe-public-access-plan).

