## Supplementary Material for "Climatic clustering and longitudinal analysis with impacts on food, bioenergy, and pandemics"

### Supporting information

**S1 Appendix. Systems used.**

1. Summit is currently the fastest computing system in the United States. It is an IBM AC922 system composed of 4662 nodes, recently upgraded from 4608, each equipped with two 22-core IBM POWER9 processors and six Nvidia Volta V100 GPUs each connected to a POWER9 by NVLink-2 with 100 GB/sec bidirectional peak performance. V100 GPU peak double precision performance is approximately 7 TF, theoretical peak mixed precision performance (FP16 accumulated to FP32) is approximately 125 TeraOps/sec, and peak memory bandwidth of 900 GB/sec. Each GPU is power capped at 300W, so the peak achievable mixed precision rate is roughly 113 TeraOps. Each node contains 512 GB of main memory, while each GPU contains 16 GB HBM2 memory. The nodes are connected by a Mellanox Infiniband fat tree interconnect with adaptive routing. Each node is equipped with a 1.6 TB NVMe burst buffer device. Summit is connected to the GPFS file system Alpine. Software versions used are GCC 6.4.0, Spectrum MPI 10.3.1.2-20200121, CUDA 10.1.243 and cuBLAS 10.2.1.243 for GEMM calculations. The `jsrun` tool is used for application launch, with six MPI ranks per node each with 1 GPU and 7 OpenMP threads mapped to CPU cores. Environment variables `PAMI_IBV_ENABLE_DCT=1 PAMI_ENABLE_STRIPING=1 PAMI_IBV_ADAPTER_AFFINITY=0 PAMI_IBV_QP_SERVICE_LEVEL=8 PAMI_IBV_ENABLE_OOO_AR=1` are set to use adaptive routing and to limit per-node storage needed for MPI buffers.

2. JUWELS Booster is currently the fastest computing system in Europe. It is a Bull Sequana XH2000 system composed of 936 nodes, each equipped with two 24-core AMD EPYC 7402 processors and four Nvidia Ampere A100 GPUs. Approximately 840 nodes were available for use in this work. Nvidia specifications for GPU peak performance report peak double precision rate at 9.7 TF, peak double precision tensor core rate at 19.5 TF, and peak tensor core 1-bit GEMM rate at 4992 TeraOps [1]. Each GPU is power capped at 400W, and the peak 1-bit GEMM rate achievable over a sustained run time using the CUTLASS library is roughly 4050 TeraOps; this implies the theoretical peak for the 840 node set used here is 16.77 1-bit GEMM ExaOps and achievable peak roughly 13.61 1-bit GEMM ExaOps. Each node contains 512 GB of main memory, while each GPU contains 40 GB HBM2 memory. The nodes are connected by Mellanox Infiniband Dragonfly+ network. A GPFS file system is used for parallel I/O. Software versions used are GCC 9.3, OpenMPI 4.1.0rc1, CUB 1.8.0, CUDA 11.2 and CUTLASS 2.3.0 for GEMM calculations. The SLURM srun command is used for job launch, with a launch command of the form `env OMP_PROC_BIND=spread OMP_PLACES=sockets OMP_NUM_THREADS=24 srun -n $num_ranks -c 24 -G $num_ranks --cpu-bind=mask_ldoms:0xc,0x3,0xc0,0x30`.

**S1 Figure. GPU utilization.** JUWELS Booster GPU usage for the global 3-way DUO vector comparison of the 500,710-coordinate dataset, with each geolocation vector containing 504,000 binary elements.

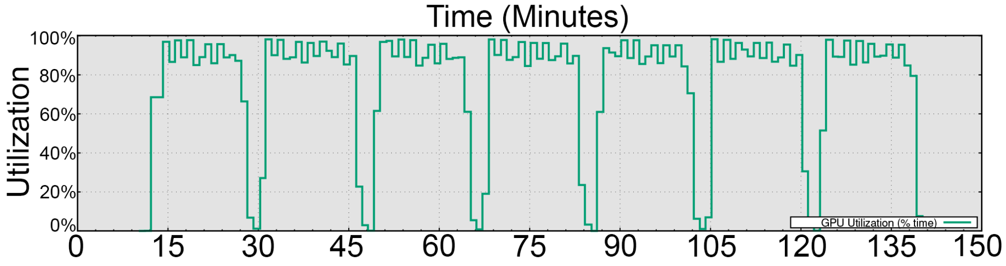


**S2 Figure. Weak scaling study.** Weak scaling performance on JUWELS Booster.

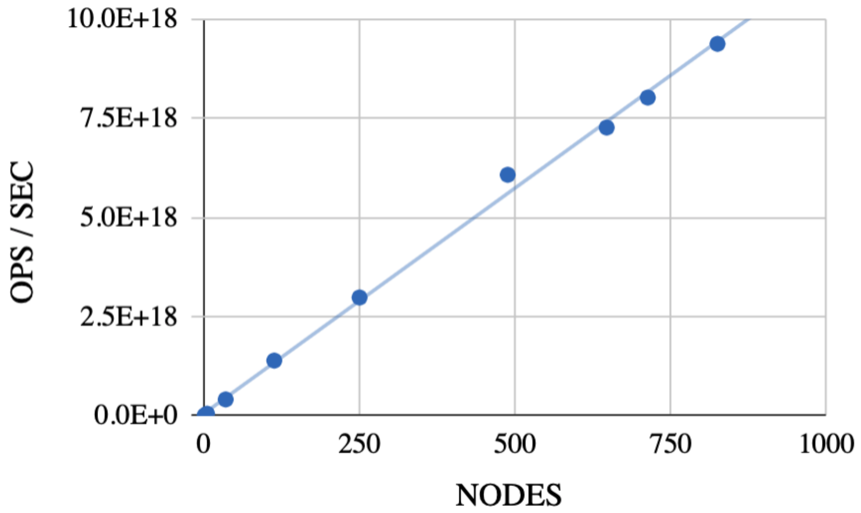

**S3 Figure. Example coordinate grid.** Example view of a coordinate grid over the United States before eliminating non-land coordinates. The grid resolution is exaggerated to facilitate visualization. Actual density used in this work is significantly greater.

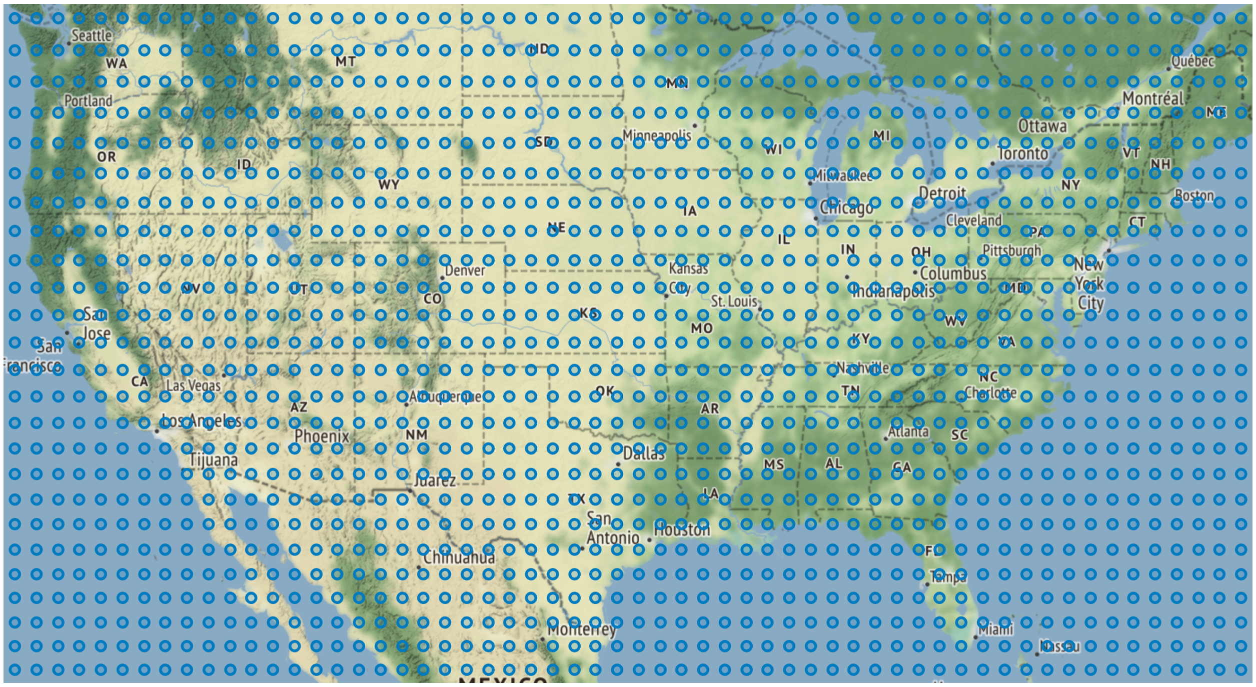

**S4 Figure. Pairwise correlation analysis.** Snapshot of Pearson's correlation coefficients between 14 climate layers from 2018 using 10,000 randomly selected global geolocations. See S1 Table for the variable acronyms.

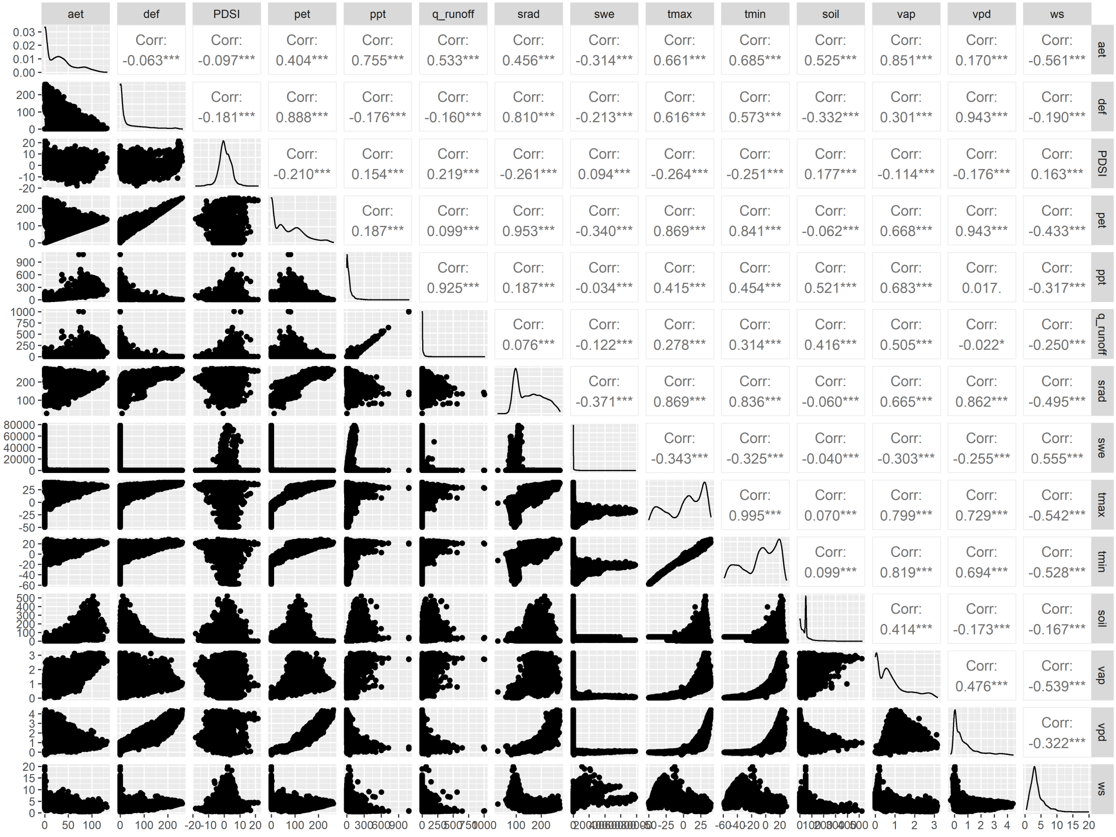

**S1 Table. Climate layer acronyms.**

| Acronym | Climate layer |
|---|---|
| aet | Actual Evapotranspiration |
| def | Climate Water Deficit |
| pet | Potential Evapotranspiration |
| ppt | Precipitation |
| q | Runoff |
| soil | Soil Moisture |
| srad | Downward Surface Shortwave Radiation |
| swe | Snow Water Equivalent |
| tmax | Max Temperature |
| tmin | Min Temperature |
| vap | Vapor Pressure |
| ws | Wind Speed |
| vpd | Vapor Pressure Deficit |
| PDSI | Palmer Drought Severity Index |

**S5 Figure. Sparse grid sub-sampling scheme.** Partial view of a sparse grid of coordinates centralized in North America. The resolution of the sparse grid points in blue are exaggerated to facilitate visualization.

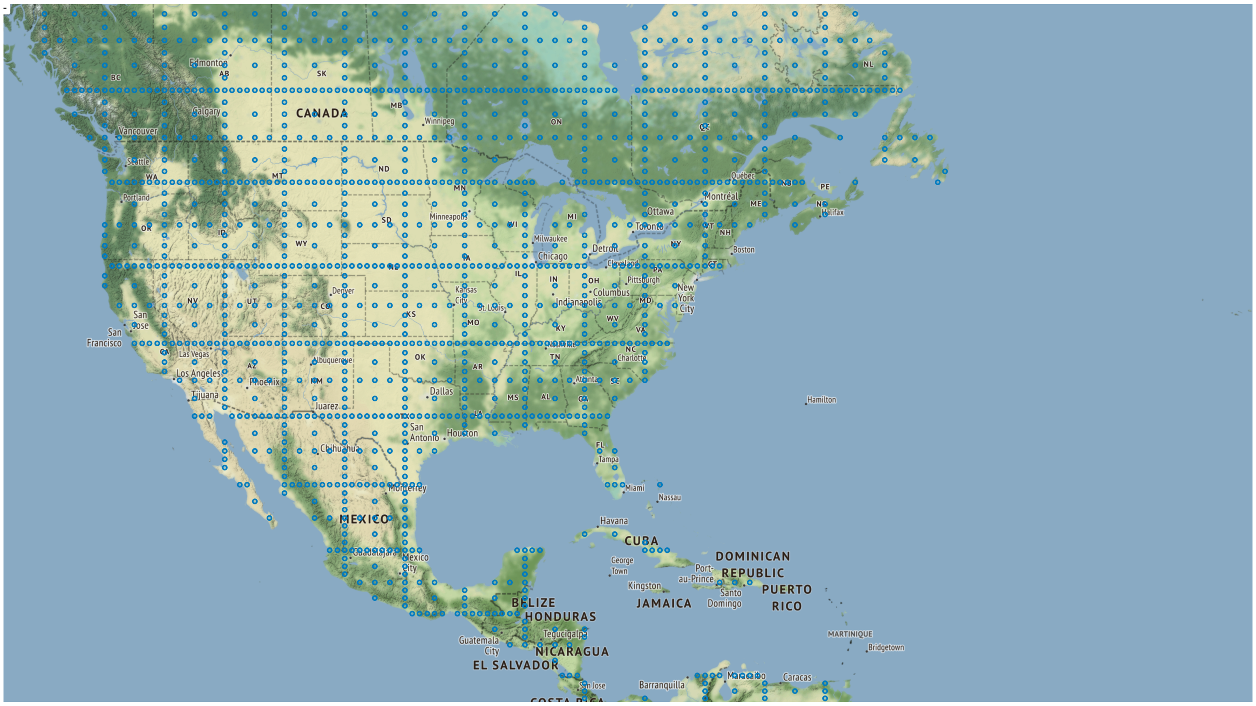
